# Leakage-controlled benchmarking reveals generalization limits of deep learning for protein–ligand binding affinity prediction

**DOI:** 10.64898/2026.09.17.752433

**Authors:** Lyuwei Wang, Jianlin Cheng

## Abstract

To address widespread data leakage and inconsistent evaluation in protein–ligand affinity prediction, we introduce PLABench, a leakage-controlled and target-centric benchmarking framework that enables standardized comparison across sequence- and structure-based models. We benchmark nine deep learning methods across blind CASP16 targets, leakage-controlled ChEMBL35 sets, and established Davis and KIBA datasets under rigorous data split settings, standardizing structural input via AlphaFold3 to ensure fair comparison. Although pretrained structure-based models achieve the highest overall accuracy, they show severe target-dependent variability, and increasing structural fidelity from predicted to experimental conformations yields no consistent gains. Meanwhile, sequence-based approaches surpass some structure-based methods on select targets, and protein family-level evaluations reveal uneven performance across families and substantial inter-model complementarity obscured by aggregate metrics. These findings demonstrate that training scale and structural input alone cannot guarantee cross-target generalization, highlighting the need for context-aware interaction modeling. PLABench provides an extensible open-source platform to facilitate these developments.

## Introduction

Accurately characterizing molecular interactions is fundamental to computational drug discovery. Protein–ligand binding affinity (PLA) quantifies the strength of these interactions and serves as a key determinant of the biological activity and mechanism of action of drug candidates. Accurate PLA prediction can accelerate compound screening, reduce experimental costs, and facilitate drug development.^1, 2^ Traditionally, the determination of PLA relies on experimental platforms such as protein microarrays and label-free detection techniques like Surface Plasmon Resonance (SPR).^3^ Although these methods can provide reliable data, they suffer from low throughput and high sensitivity to experimental conditions, limiting their application in large-scale drug screening. Therefore, there is a significant need to develop computational PLA prediction methods. In the last several years, deep learning models have emerged as core tools in PLA prediction and computer-aided drug design due to their powerful feature learning capabilities.

Contemporary deep learning-based PLA prediction methods can be broadly classified into sequence-based and structure-based approaches. The former typically utilizes 1D protein sequences and ligand SMILES strings as input, offering rapid inference and broad applicability, albeit at the cost of neglecting critical spatial information.^4, 5^ Conversely, structure-based approaches have emerged as the dominant paradigm in recent years, as their ability to explicitly integrate 3D structural data enables a more direct modeling of spatial intermolecular interactions.^6, 7^

However, despite the rapid proliferation of prediction models, the field still lacks a standardized benchmarking framework to quantify the actual progress of these methods.^8^ Existing evaluations suffer from the following three primary limitations:

First, modality discrepancy and incomparability: Sequence-based methods are typically evaluated on large-scale sequence-based datasets, such as Davis and KIBA. However, because these datasets lack 3D spatial conformations of protein targets, structure-based methods cannot be deployed on them. This disconnect creates a barrier, making it difficult to fairly compare the performance and generalizability across the two major prediction paradigms.

Second, severe data leakage: As newly developed deep learning models leverage increasingly massive datasets for training, traditional test sets like CASF^9, 10^ and CSAR^11^ face an escalating crisis of data leakage. Many modern models have implicitly or explicitly “seen” these test samples during their training phase, leading to overestimated performance and severely biased conclusions regarding their true predictive capabilities.

Third, lack of target-centric evaluation: Standard benchmarks (e.g., CASF and CSAR) are primarily designed to evaluate absolute affinity across a broad, highly diverse set of protein–ligand complexes. Consequently, they fail to assess a model’s ability to accurately rank the relative or absolute affinities of different compounds binding to the same target protein. However, this target-centric ranking is fundamentally closer to real-world drug design scenarios (e.g., lead optimization), where distinguishing subtle affinity differences among congeneric ligands is critical.

Here, we systematically investigate the generalization capabilities and limitations of current deep learning approaches for protein–ligand binding affinity prediction through PLABench, a standardized evaluation framework designed to minimize data leakage and enable direct comparison across model paradigms. We evaluate nine recent sequence- and structure-based models using target-centric CASP16 and ChEMBL35 datasets, together with established affinity datasets (Davis and KIBA). To enable comparison under common structural conditions, we generate AlphaFold3 structures for test cases lacking experimental structures and retrain sequence-based models on a common PDBbind training corpus. Our analyses reveal that although massively pretrained structure-based models achieve the strongest aggregate performance, their advantages vary substantially across protein families and disappear for some targets. Interestingly, progressively improving protein–ligand structural accuracy up to experimental crystal structures does not consistently improve affinity prediction, indicating that current structure-based models do not fully exploit the additional molecular information provided by more accurate binding conformations. We further identify target classes in which current models systematically struggle and show substantial complementarity between model paradigms. Together, these findings reveal limitations that are obscured by conventional aggregate benchmarks and establish a leakage-controlled framework for developing affinity models with improved cross-target generalization.

## Results

The evaluation workflow of PLABench is depicted in Fig. 1. The performance of nine deep learning PLA prediction methods (six structure-based: Boltz2,^6^ FLOWR.ROOT,^12^ Graph_RG,^13^ LCDD-team,^14^ FlowDock,^7^ MFE;^15^ three sequence-based: DeepDTA,^4^ LLF,^16^ MixingDTA^5^) on four test datasets is assessed in PLABench. The deep learning architecture, original training and test datasets, and inputs of the nine methods are briefly described in Table 1.

**Fig. 1.**
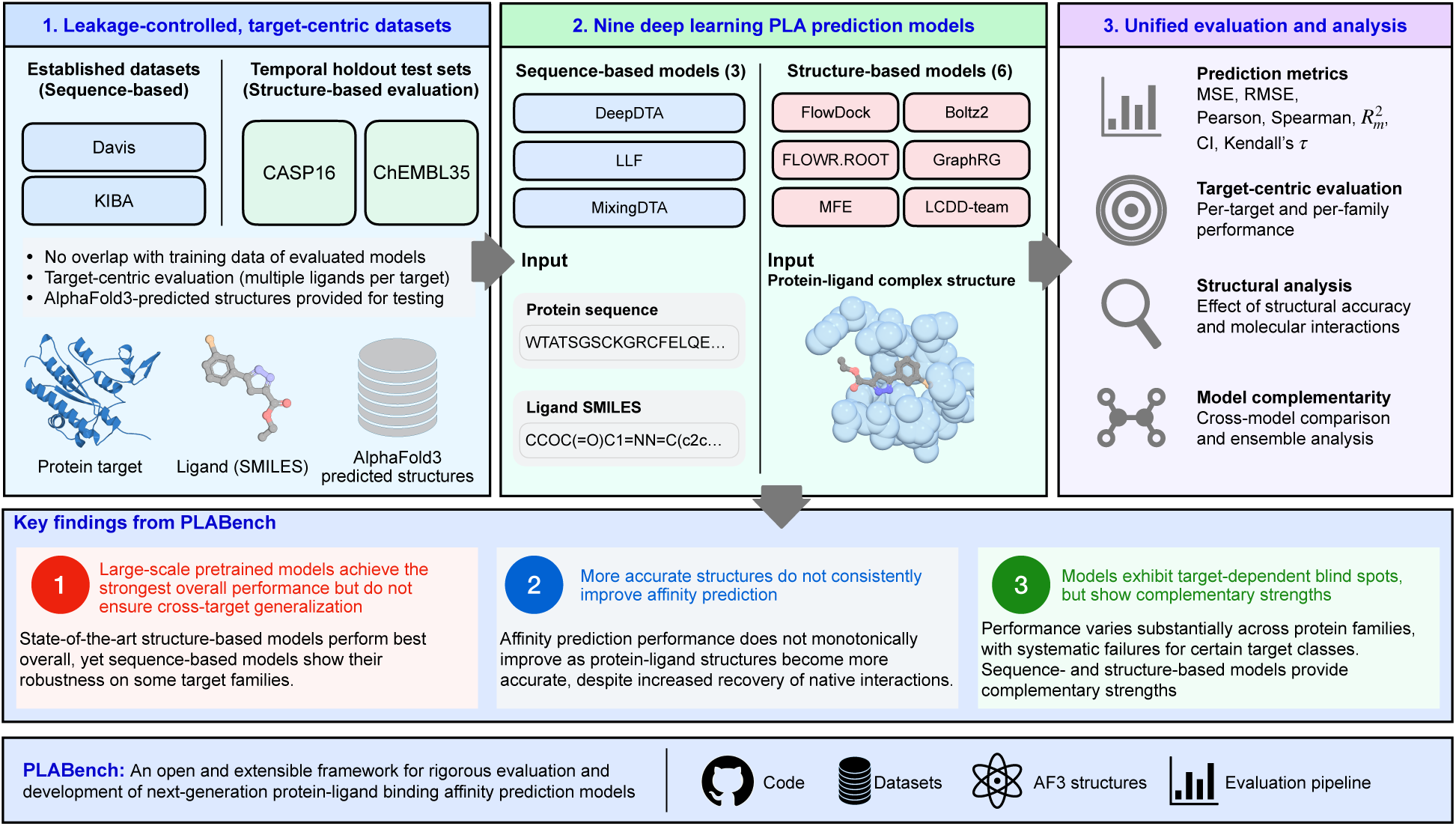
Overview of PLABench and its major findings. PLABench provides a leakage-controlled framework for evaluating nine protein–ligand binding affinity prediction methods, including three sequence-based models (DeepDTA, LLF, and MixingDTA) and six structure-based models (Boltz2, FLOWR.ROOT, Graph_RG, LCDD-team, FlowDock, and MFE). The framework couples established affinity benchmarks (Davis, KIBA) with prospective (CASP16) and leakage-controlled (ChEMBL35) test sets, supplying AlphaFold3-predicted conformations when experimental complexes are unavailable. Models are evaluated using prediction-error, correlation, and ranking metrics, together with target-centric, structural, and protein-family analyses. The evaluation reveals key strengths and limitations: large-scale pretrained models achieve the strongest overall performance but do not ensure cross-target generalization; increasing structural accuracy does not consistently improve affinity prediction; and different models exhibit target-dependent blind spots and complementary strengths. PLABench provides an open and extensible framework for evaluating future protein–ligand affinity prediction methods.

**Table 1.** An overview of 9 sequence-based and structure-based PLA prediction models.

| Category | Name | Year | Deep Learning Architecture | Train Set | Test Set | Input Type |
| --- | --- | --- | --- | --- | --- | --- |
| Sequence-based | DeepDTA<sup>4</sup> | 2018 | Two-stream CNN | 5/6 of Davis & KIBA, 5-fold cross validation | 1/6 of Davis & KIBA | Seq + SMILES |
|  | LLF<sup>16</sup> | 2025 | Multi-Stream GCN, CNN; Keep Local features | Davis, KIBA, 5-fold cross validation | Davis, KIBA, 5-fold cross validation | Seq + SMILES |
|  | MixingDTA<sup>5</sup> | 2025 | MolFormer<sup>17</sup> & ESM3 feature extraction;<sup>18</sup> Transformer; Meta-Ensemble | Davis, KIBA, 4/5 warm start, 4/5 cold-drug/target split | Davis, KIBA, 1/5 warm start, 1/5 cold-drug/target split | Seq + SMILES |
| Structure-based | Graph_RG<sup>13</sup> | 2025 | Two-stream Graph Transformer; Ligand pose-free | PDBbind v2020, 20k | CASP16 | Protein PDB + SMILES |
|  | Boltz2<sup>6</sup> | 2025 | MSA, PairFormer,<sup>19</sup> Diffusion, Hybrid Prediction, AlphaFold2<sup>20</sup> and Boltz-1 Distillation | ChEMBL, BindingDB, PubChem ~1.21M | FEP+, CASP16, MF-PCBA, Internal Industry Assays | Seq + SMILES |
|  | FlowDock<sup>7</sup> | 2025 | ESMFold,<sup>21</sup> NeuralPLexer,<sup>22</sup> Conditional Flow Matching, Hybrid Prediction | PDBbind v2020 (before 2019, 17,743) BindingMOAD (46,567) | PDBbind (after 2019, 363) CASP16 | Protein PDB + Ligand SDF |
|  | FLOWR.ROOT<sup>12</sup> | 2025 | SE(3)-equivariant Flow Matching, Joint Structure-Affinity-Confidence Heads | SPINDR, Plinder, BindingMOAD, SAIR, HiQBind, BindingNet, KIBA-3D, DAVIS-3D, Kinodata-3D ~2.5M Complexes | GEOM-DRUGS, CROSS-DOCKED2020, SPINDR, Schrödinger FEP+, OpenFE, Internal Assays | Protein PDB + Ligand SDF |
|  | MFE<sup>15</sup> | 2024 | Multimodal Extraction, 3D CNN, GVP-GNN,<sup>23</sup> ProtTrans,<sup>24</sup> Transformer Fusion | PDBbind v2016, General + Refined (except Core Set) | PDBbind v2016 Core Set (285) | Protein PDB + Ligand SDF |
|  | LCDD-team<sup>14</sup> | 2025 | 3-stream GPS Graph Transformer | Unknown | CASP16 | Protein PDB + Ligand SDF |

*Abbreviations:* CNN, Convolutional Neural Network; GCN, Graph Convolutional Network; GNN, Graph Neural Network; GVP, Geometric Vector Perceptron; MSA, Multiple Sequence Alignment.

As shown in Table 1, these PLA prediction methods were originally trained and tested on different datasets. To ensure a fair comparison, the three sequence-based methods that can be easily retrained were re-trained on the PDBbind v2020 refined dataset^25^ before they were evaluated in PLABench. In contrast, because exact training recipes and end-to-end training scripts are not publicly available for most structure-based methods, we evaluated them using their official pre-trained checkpoints to ensure their optimal reported performance, with potential training contamination systematically controlled through chemical-similarity filtering and cross-corpus auditing.

The subsequent evaluations are organized progressively to assess the generalization capability of the models: beginning with classic sequence-based PLA test datasets (Davis^26^ and KIBA^27^), then testing with the highly challenging real-world CASP16 competition dataset,^28^ and finally benchmarking with the leakage-controlled, target-centric ChEMBL35 dataset.^29^

To ensure a multi-dimensional evaluation, we measure the prediction error (MSE, RMSE) between true and predicted PLA, correlation (Pearson’s *r*, Spearman’s *ρ*, *r*^2^_m_), and target-specific ranking capability (CI, Kendall’s *τ*) of each method, with the formal definitions of these metrics detailed in Methods (‘Evaluation metrics’).

### Sequence-based affinity models show substantial generalization loss on unseen targets and ligands

The Davis and KIBA datasets serve as standard benchmarks for sequence-based methods. To ensure a fair comparison among the three sequence-based models, we retrained them using the specific warm split and cold split configurations established by MixingDTA.^5^

Specifically, the warm split involves a random partition of the dataset (typically evaluated via 5-fold cross-validation) to verify the models’ basic pattern recognition capabilities. Conversely, the cold-target split ensures that the protein targets in the test set are entirely unseen during the training and validation phases, while the cold-drug split applies the same strict non-overlapping criterion to the ligand molecules. These cold splits evaluate generalization to unseen targets and chemotypes, reflecting conditions typical of prospective drug discovery. The results are shown in Table 2.

**Table 2.**
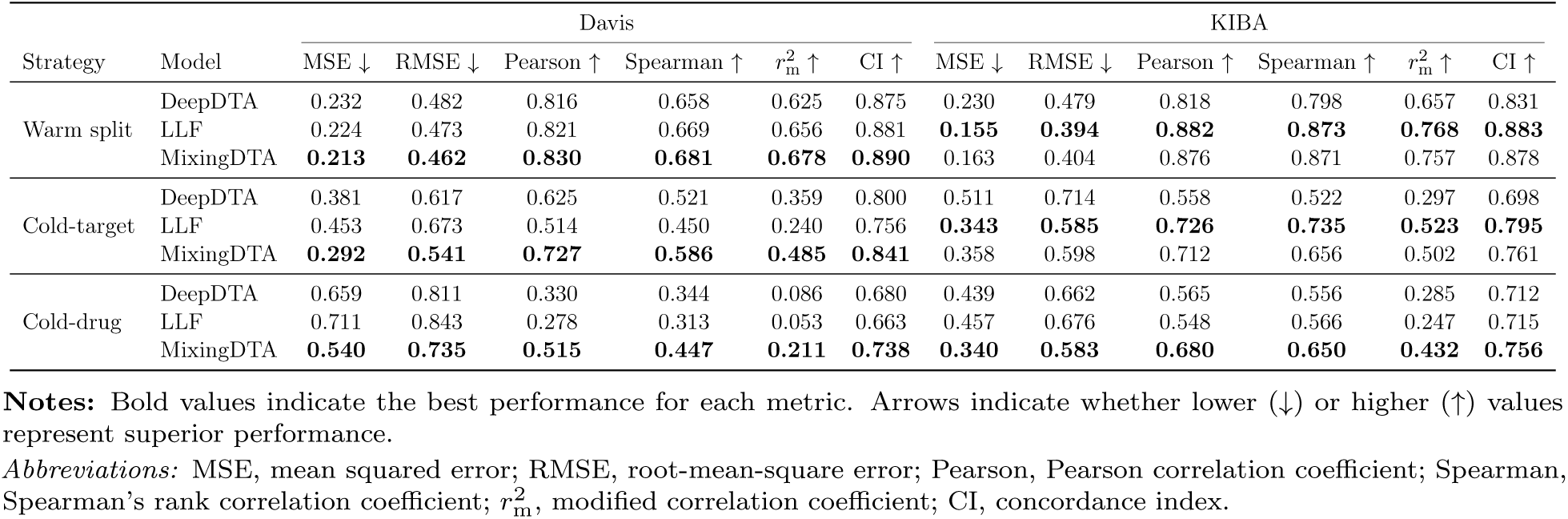
Performance comparison of different models on Davis and KIBA datasets under various splitting strategies.

| Strategy | Model | Davis |  |  |  |  |  | KIBA |  |  |  |  |  |
| --- | --- | --- | --- | --- | --- | --- | --- | --- | --- | --- | --- | --- | --- |
| | | MSE ↓ | RMSE ↓ | Pearson ↑ | Spearman ↑ | $r_m^2$ ↑ | CI ↑ | MSE ↓ | RMSE ↓ | Pearson ↑ | Spearman ↑ | $r_m^2$ ↑ | CI ↑ |
| Warm split | DeepDTA | 0.232 | 0.482 | 0.816 | 0.658 | 0.625 | 0.875 | 0.230 | 0.479 | 0.818 | 0.798 | 0.657 | 0.831 |
|  | LLF | 0.224 | 0.473 | 0.821 | 0.669 | 0.656 | 0.881 | **0.155** | **0.394** | **0.882** | **0.873** | **0.768** | **0.883** |
|  | MixingDTA | **0.213** | **0.462** | **0.830** | **0.681** | **0.678** | **0.890** | 0.163 | 0.404 | 0.876 | 0.871 | 0.757 | 0.878 |
| Cold-target | DeepDTA | 0.381 | 0.617 | 0.625 | 0.521 | 0.359 | 0.800 | 0.511 | 0.714 | 0.558 | 0.522 | 0.297 | 0.698 |
|  | LLF | 0.453 | 0.673 | 0.514 | 0.450 | 0.240 | 0.756 | **0.343** | **0.585** | **0.726** | **0.735** | **0.523** | **0.795** |
|  | MixingDTA | **0.292** | **0.541** | **0.727** | **0.586** | **0.485** | **0.841** | 0.358 | 0.598 | 0.712 | 0.656 | 0.502 | 0.761 |
| Cold-drug | DeepDTA | 0.659 | 0.811 | 0.330 | 0.344 | 0.086 | 0.680 | 0.439 | 0.662 | 0.565 | 0.556 | 0.285 | 0.712 |
|  | LLF | 0.711 | 0.843 | 0.278 | 0.313 | 0.053 | 0.663 | 0.457 | 0.676 | 0.548 | 0.566 | 0.247 | 0.715 |
|  | MixingDTA | **0.540** | **0.735** | **0.515** | **0.447** | **0.211** | **0.738** | **0.340** | **0.583** | **0.680** | **0.650** | **0.432** | **0.756** |

**Notes:** Bold values indicate the best performance for each metric. Arrows indicate whether lower (↓) or higher (↑) values represent superior performance.
*Abbreviations:* MSE, mean squared error; RMSE, root-mean-square error; Pearson, Pearson correlation coefficient; Spearman, Spearman’s rank correlation coefficient; $r_m^2$ , modified correlation coefficient; CI, concordance index.

MixingDTA employs a data augmentation strategy known as guilt-by-association (GBA) Mixup,^5^ which enables it to achieve leading performance across the majority of evaluation settings on both the Davis and KIBA datasets. Nevertheless, LLF^16^ achieves the strongest performance on KIBA under both the warm and cold-target splits. Notably, despite being introduced in 2018, DeepDTA^4^ remains surprisingly competitive against its 2025 counterpart, LLF, in some experiments, outperforming it in both the cold-drug and cold-target evaluations on the Davis dataset.

For all three sequence-based models, their performance in the cold-target or cold-drug evaluation scheme is generally worse than in the warm split schedule. And in many cases, the performance deterioration is substantial, indicating some limitation in generalizing to new targets or drugs.

### State-of-the-art affinity models remain limited in ranking closely related high-affinity ligands

The CASP16 (2024) dataset comprises two clinically relevant targets from active pharmaceutical discovery programs, Chymase (series L1000, *n* = 17 ligands) and Autotaxin (series L3000, *n* = 123 ligands). All nine sequence- and structure-based methods were evaluated off-the-shelf without any target-specific fine-tuning.

To quantify the extent of data isolation across the evaluated models, we performed a comprehensive cross-corpus audit against the training datasets of all evaluated models. This screen revealed that a subset of CASP16 ligands existed in historical bioactivity databases prior to the challenge: Boltz2’s training corpora (ChEMBL34 and BindingDB) contain 1 of the 17 L1000 ligands and 19 of the 123 L3000 ligands, while FLOWR.ROOT’s pretraining sets (SAIR and BindingNet) contain 1 and 14 pairs, respectively (with no overlap detected for the remaining models). Although the historical bioactivity values deviate from the blinded CASP16 measurements by a median absolute difference of 0.35 log units (consistent with expected inter-assay experimental variability), we retain the complete series in our primary analysis to maintain direct comparability with the official CASP16 evaluation standard. Excluding these overlapping pairs does not alter the relative rankings of Boltz2 and FLOWR.ROOT, which consistently lead the benchmark across all evaluation metrics (complete de-duplicated performance metrics are provided in Supplementary Note 3).

As shown in Fig. 2, Boltz2 achieves the highest overall performance in terms of Kendall rank correlation coefficient, indicating stronger generalization to these blind-test series. While Graph_RG^13^ secured the top rank in the official CASP16 competition in 2024, it falls behind Boltz2^6^ and FLOWR.ROOT^12^ that were developed after the conclusion of CASP16 in our benchmark. This discrepancy reflects a progress: the massive scale of training data utilized by Boltz2 and FLOWR.ROOT (trained on approximately 1.21M and 2.5M complexes, roughly 60-fold and 125-fold larger than Graph_RG’s *∼*20k dataset, respectively) may have helped improve prediction accuracy.

**Fig. 2.**
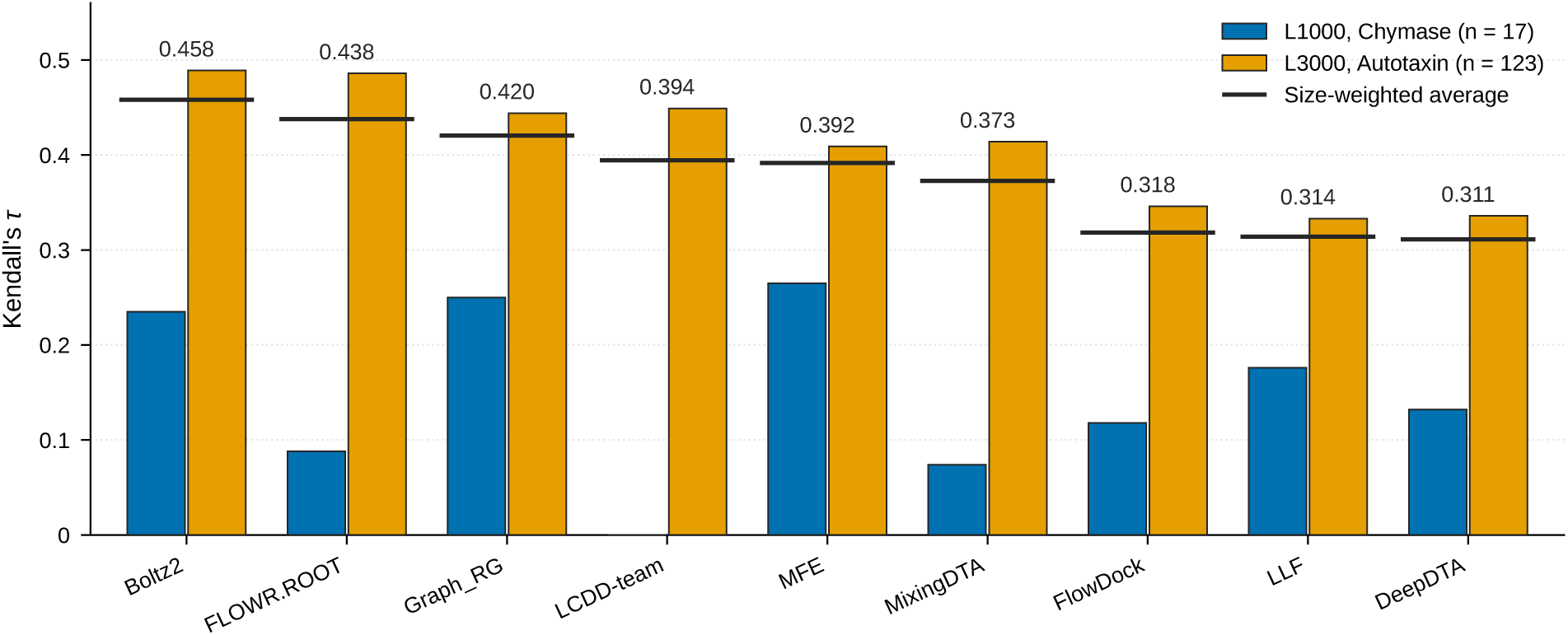
Kendall’s *τ* performance across CASP16 data series. Grouped bars display the individual Kendall’s *τ* scores for the Chymase series L1000 (*n* = 17 complexes, blue) and Autotaxin series L3000 (*n* = 123 complexes, orange). The black horizontal lines denote the final weighted average *τ* score for each model, with precise numerical values annotated above. Models are sorted in descending order based on this weighted average. Each *τ* is a single value computed over all ligands of a series, so no error bars are shown. The L1000 bar for LCDD-team is not visible because its *τ* is 0.000.

Furthermore, it is evident that structure-based models generally outperform sequence-based models (i.e., LLF, MixingDTA, and DeepDTA) in this test scenario. To ensure a fair comparison of structure-based methods and to address the issue that some of them (e.g., FlowDock^7^ and LCDD-team^14^) do not predict protein structures themselves and require structural inputs, we standardized the evaluation pipeline by supplying the same AlphaFold3-predicted structures to all structure-based models. Notably, this arrangement did not degrade model performance (shown in Supplementary Note 2), indicating that AlphaFold3 conformations serve as adequate inputs for downstream affinity prediction.

A distinguishing characteristic of the CASP16 PLA dataset is its concentrated distribution of high binding affinities (L1000: p*K*_d_ = 7.69 *±* 0.60; L3000: p*K*_d_ = 6.94 *±* 1.13). The narrow variance poses a rigorous challenge, revealing an intriguing paradox in Table 3: models that excel in regression accuracy (RMSE) frequently diverge from those that demonstrate the stronger ranking capabilities. For example, in the L1000 series, while Boltz2 achieves the lowest prediction error (RMSE 0.660), MFE dominates the ranking metrics (Spearman 0.485, CI 0.632) despite exhibiting a significantly higher error (RMSE 1.235). In contrast, in the L3000 series, FLOWR.ROOT yields the lowest RMSE (0.854), but Boltz2 leads in most of correlation and ranking metrics. This phenomenon suggests that achieving a low RMSE in such constrained regimes might merely result from models safely “learning the mean” of the distribution, rather than distinguishing the relative difference in the binding affinities of different ligands.

**Table 3.** Performance comparison of different models on the CASP16 dataset (L1000 and L3000 ligand series).

| Model | L1000 |  |  |  |  |  | L3000 |  |  |  |  |  |
| --- | --- | --- | --- | --- | --- | --- | --- | --- | --- | --- | --- | --- |
| | MSE ↓ | RMSE ↓ | Pearson ↑ | Spearman ↑ | $r_m^2$ ↑ | CI ↑ | MSE ↓ | RMSE ↓ | Pearson ↑ | Spearman ↑ | $r_m^2$ ↑ | CI ↑ |
| Boltz2 | **0.436** | **0.660** | **0.544** | 0.382 | **0.068** | 0.618 | 1.034 | 1.017 | **0.670** | **0.670** | 0.356 | **0.745** |
| FlowDock | 2.559 | 1.600 | 0.142 | 0.140 | -0.006 | 0.559 | 1.316 | 1.147 | 0.462 | 0.490 | 0.178 | 0.674 |
| FLOWR.ROOT | 1.382 | 1.176 | 0.222 | 0.162 | 0.030 | 0.544 | **0.730** | **0.854** | 0.661 | 0.669 | **0.432** | 0.744 |
| Graph_RG | 1.306 | 1.143 | 0.406 | 0.404 | -0.020 | 0.625 | 1.075 | 1.037 | 0.576 | 0.632 | 0.312 | 0.723 |
| LCDD-team | 1.139 | 1.067 | 0.054 | 0.022 | 0.000 | 0.500 | 0.843 | 0.918 | 0.615 | 0.628 | 0.339 | 0.725 |
| MFE | 1.525 | 1.235 | 0.332 | **0.485** | 0.037 | **0.632** | 1.318 | 1.148 | 0.613 | 0.574 | 0.345 | 0.705 |
| DeepDTA | 1.018 | 1.009 | -0.195 | 0.201 | -0.008 | 0.566 | 1.582 | 1.258 | 0.485 | 0.475 | 0.200 | 0.668 |
| LLF | 0.536 | 0.732 | 0.181 | 0.279 | 0.020 | 0.588 | 3.385 | 1.840 | 0.484 | 0.467 | 0.232 | 0.667 |
| MixingDTA | 2.876 | 1.696 | -0.085 | 0.113 | -0.004 | 0.537 | 0.874 | 0.935 | 0.597 | 0.587 | 0.331 | 0.708 |

**Notes:** Bold values indicate the best performance for each metric. Arrows indicate whether lower (↓) or higher (↑) values represent superior performance.
*Abbreviations:* MSE, mean squared error; RMSE, root-mean-square error; Pearson, Pearson correlation coefficient; Spearman, Spearman’s rank correlation coefficient; $r_m^2$ , modified correlation coefficient; CI, concordance index.

Indeed, the absolute ranking performance of current state-of-the-art models is moderate: the best Kendall’s *τ* achieved by Boltz2 (*∼* 0.46) falls significantly below the theoretical upper limit of approximately 0.73.^28^

### Increasing structural accuracy does not consistently improve affinity prediction

Previous CASP16 assessment showed that providing experimental crystal structures did not improve affinity prediction over AlphaFold3-predicted conformations.^28^ However, it remains unclear whether this apparent structural insensitivity persists systematically across different levels of structural accuracy and whether increasingly accurate structures provide molecular interactions that current affinity models fail to exploit. To investigate this question, we designed a systematically controlled experiment on the CASP16 L3000 series across a four-tier ladder of input pose accuracy (Fig. 3): (1) the predicted target–ligand complex structure of Boltz2 unguided co-folding (median ligand RMSD 6.96 Å), (2) the predicted structure of Boltz2 guided by the experimental receptor template with an enforced pocket constraint (3.72 Å), (3) AlphaFold3 top-1 predicted complex structure (0.76 Å), and (4) the deposited crystal complex structure (0 Å). Importantly, all ligands in these evaluations bound within the same canonical orthosteric cavity, precluding off-pocket situations. The sample size is reduced from 123 to 93 because 30 complexes lack experimental input structures.

**Fig. 3.**
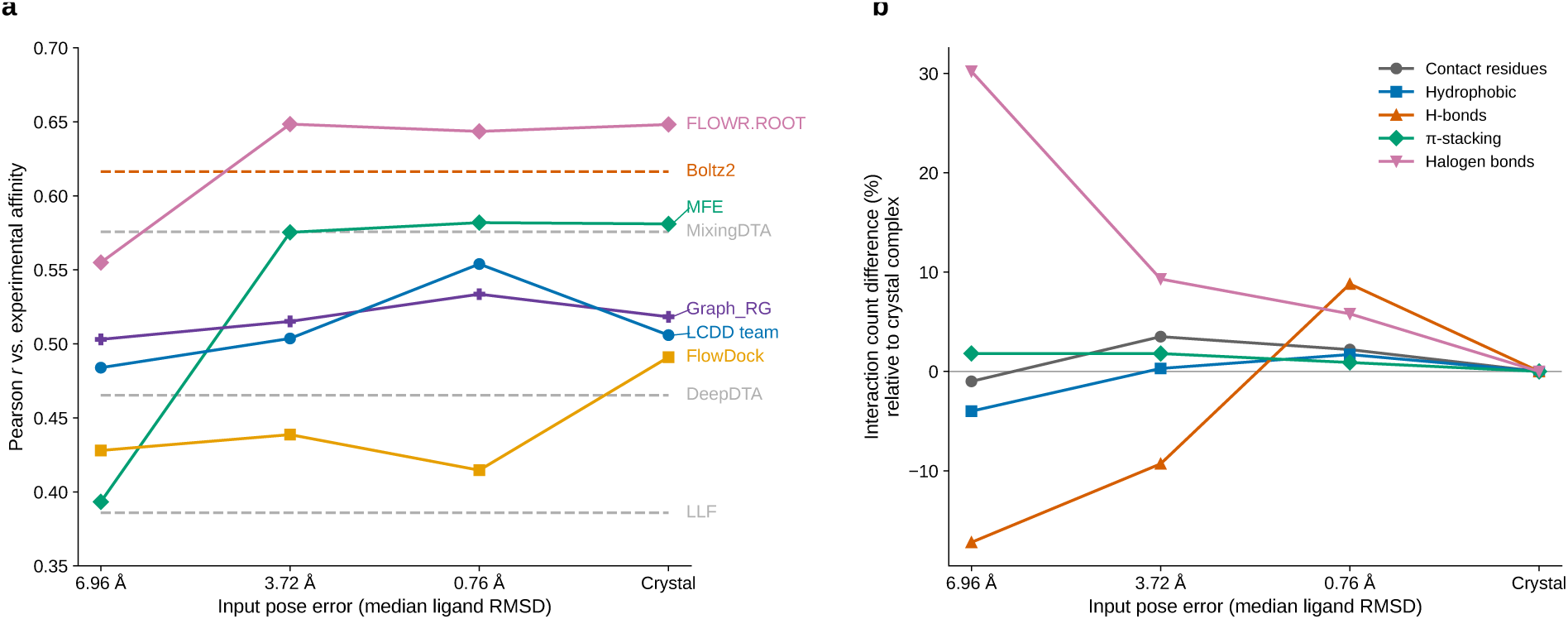
Impact of input pose accuracy in terms of RMSD on binding affinity prediction and molecular interactions. **a**, Pearson *r* against CASP16 L3000 (*n* = 93) experimental affinity for six structure-based models across four input structural conditions: (1) the predicted protein–ligand complex structure of unguided co-folding by Boltz2 (median ligand RMSD 6.96 Å), (2) the predicted structure of Boltz2 with the experimental receptor template and pocket constraint (3.72 Å), (3) the AlphaFold3 top-1 predicted complex structure (0.76 Å), and (4) the deposited crystal complex (0 Å). Boltz2 and the three sequence-based models, which do not consume the input structure, are drawn as dashed horizontal lines **b**, Percentage difference in the counts of five types of inter-atomic protein–ligand interactions relative to the experimental crystal pose (salt bridges omitted; only one occurs in the crystal complexes).

As shown in Fig. 3**a**, model sensitivity to input pose fidelity deviates from the conventional assumption that structural precision monotonically drives affinity accuracy. Among the evaluated structure-based methods, only MFE exhibited a monotonic increase in Pearson’s *r* with improved pose quality; however, the performance of some models such as FLOWR.ROOT and MFE was largely saturated early at 3.72 Å, eventually plateauing. For instance, FLOWR.ROOT, which achieved the strongest overall correlation on experimental structures, showed negligible variance across conformations within the sub-3.72 Å regime, not benefiting from near-native (0.76 Å) or crystal (0 Å) coordinates. Boltz2, along with sequence-based models, is shown as a horizontal dashed line indicating a constant value because they do not use the externally provided input structures.

One likely reason that the models fail to benefit from the highest accuracy of experimental structures is their inability to discern how individual inter-atomic interactions differentially contribute to binding affinity. To investigate this possibility, we reimplemented PLIP^30^-based interaction criteria for hydrogen-free coordinates and profiled the distribution of non-covalent interactions (NCIs) across all four conformational tiers (Fig. 3**b**). Although the recovery of native NCIs systematically increased with geometric precision, recapturing 35.4%, 52.0%, and 73.1% of the crystal interactions across the 6.96 Å, 3.72 Å, and 0.76 Å tiers, respectively, predicted poses frequently formed compensatory, non-native interactions. Consequently, the total interaction counts across these tiers reached 97.0%, 99.2%, and 103.6% of those observed in the crystal complex. Notably, in the AF3-generated poses (0.76 Å), the absolute count for every evaluated NCI class exceeded that of the native crystal structure. These patterns suggest a potential explanation for the performance plateau: current PLA prediction methods may not effectively distinguish affinity-driving interactions captured in crystal structures from secondary or incidental interactions introduced by predicted conformations.

### Large-scale pretraining improves overall generalization but does not eliminate target-dependent failures

To evaluate model generalization in a large-scale data leakage-controlled scenario, we utilized an independent test set derived from the newly released ChEMBL v35,^31^ curated by Shimizu et al.^29^ To prevent test-set contamination from the training corpora of all evaluated methods, we further enforced a chemical-similarity filter: any candidate ligand exhibiting an ECFP4 Tanimoto similarity *≥* 0.85 to any molecule present in the known training datasets of the nine models (Table 1) was strictly excluded, irrespective of the associated protein target. Retaining targets with at least 10 compounds yielded a benchmark of 4,862 complexes across 71 target proteins, providing a reliable basis for target-centric ranking analyses. The results of the nine models are reported in Table 4.

**Table 4.** Performance comparison of different models on the ChEMBL35 dataset.

| Model | MSE ↓ | RMSE ↓ | Pearson ↑ | Spearman ↑ | Kendall ↑ | $r_m^2$ ↑ | CI ↑ |
| --- | --- | --- | --- | --- | --- | --- | --- |
| Boltz2 | 1.692 | 1.248 | 0.321 | 0.302 | 0.211 | 0.098 | 0.606 |
| FlowDock | 2.216 | 1.369 | 0.188 | 0.173 | 0.120 | 0.051 | 0.560 |
| FLOWR.ROOT | **0.931** | **0.928** | **0.392** | **0.374** | **0.264** | **0.168** | **0.632** |
| Graph_RG | 1.973 | 1.340 | 0.146 | 0.128 | 0.089 | 0.035 | 0.544 |
| LCDD-team | 1.636 | 1.221 | 0.157 | 0.131 | 0.091 | 0.038 | 0.545 |
| MFE | 1.626 | 1.212 | 0.177 | 0.162 | 0.111 | 0.048 | 0.556 |
| DeepDTA | 2.165 | 1.393 | 0.153 | 0.151 | 0.103 | 0.045 | 0.551 |
| LLF | 2.503 | 1.462 | 0.212 | 0.208 | 0.144 | 0.072 | 0.572 |
| MixingDTA | 1.814 | 1.272 | 0.173 | 0.162 | 0.112 | 0.048 | 0.556 |

**Notes:** Bold values indicate the best performance for each metric. Arrows indicate whether lower (↓) or higher (↑) values represent superior performance.
*Abbreviations:* MSE, mean squared error; RMSE, root-mean-square error; Pearson, Pearson correlation coefficient; Spearman, Spearman’s rank correlation coefficient; Kendall, Kendall’s rank correlation coefficient ( $\tau$ ); $r_m^2$ , modified correlation coefficient; CI, concordance index.

#### The advantage of massive data scaling and pretraining

The most notable observation is the clear performance advantage demonstrated by the two massively pretrained models, FLOWR.ROOT and Boltz2, which markedly separate themselves from all other approaches (Table 4). FLOWR.ROOT achieves the best overall performance across all evaluated metrics (Pearson *r* = 0.392, Kendall’s *τ* = 0.264, RMSE = 0.928), while Boltz2 secures the second-best performance (Pearson *r* = 0.321, Kendall’s *τ* = 0.211). Because FLOWR.ROOT and Boltz2 were trained on substantially larger datasets than the other evaluated methods, these results are consistent with an important contribution of the training-data scale to generalization, although differences in model architecture, training objectives, and data composition prevent attributing the performance gains to scale alone. The strong performance of FLOWR.ROOT may also reflect its two-stage training paradigm: leveraging 2.5 million low-fidelity data points for pre-training, followed by fine-tuning on *∼*30,000 high-fidelity points. Similarly, Boltz2 benefits from a training set of 1.21M complexes. In contrast, other structure-based methods, Graph_RG, MFE, FlowDock, and LCDD-team, predominantly trained on the PDBbind dataset, perform much worse than FLOWR.ROOT and Boltz2.

#### Competitive target-centric ligand ranking achieved by the sequence-based model LLF

While massively pre-trained structure-based models achieve the best overall performance, the sequence-based model LLF demonstrates competitive ranking capabilities. As shown in Table 4, although LLF exhibits higher absolute prediction errors (e.g., MSE of 2.503 and RMSE of 1.462) compared to most structure-based methods, it surprisingly outperforms several structure-based models (such as Graph_RG, MFE, FlowDock, and LCDD-team) across all correlation and ranking metrics (Pearson’s *r*, Spearman’s *ρ*, Kendall’s *τ*, and CI).

This divergence could not be attributed to the input distribution shift induced by AlphaFold3-predicted poses according to our findings in Fig. 3**a**. The suboptimal performance of structure-based models on ChEMBL35 therefore reflects intrinsic limitations within their affinity scoring heads, rather than deficiencies in input structural fidelity.

#### Affinity prediction models exhibit systematic protein-family-dependent blind spots

To evaluate class-dependent model behaviors, we stratified the benchmark by protein family across 5,002 unique complexes (4,862 from ChEMBL35 alongside 140 from the prospective CASP16 Chymase and Autotaxin series). Accounting for the bifunctional enzyme CD38 (12 ligands dual-assigned to both Transferases and Hydrolases), this yields 5,014 family-level evaluation pairs. As visualized in the heatmap (Fig. 4), this analysis reveals class-conditional disparities across different model architectures, highlighting specific blind spots among the models.

**Fig. 4.**
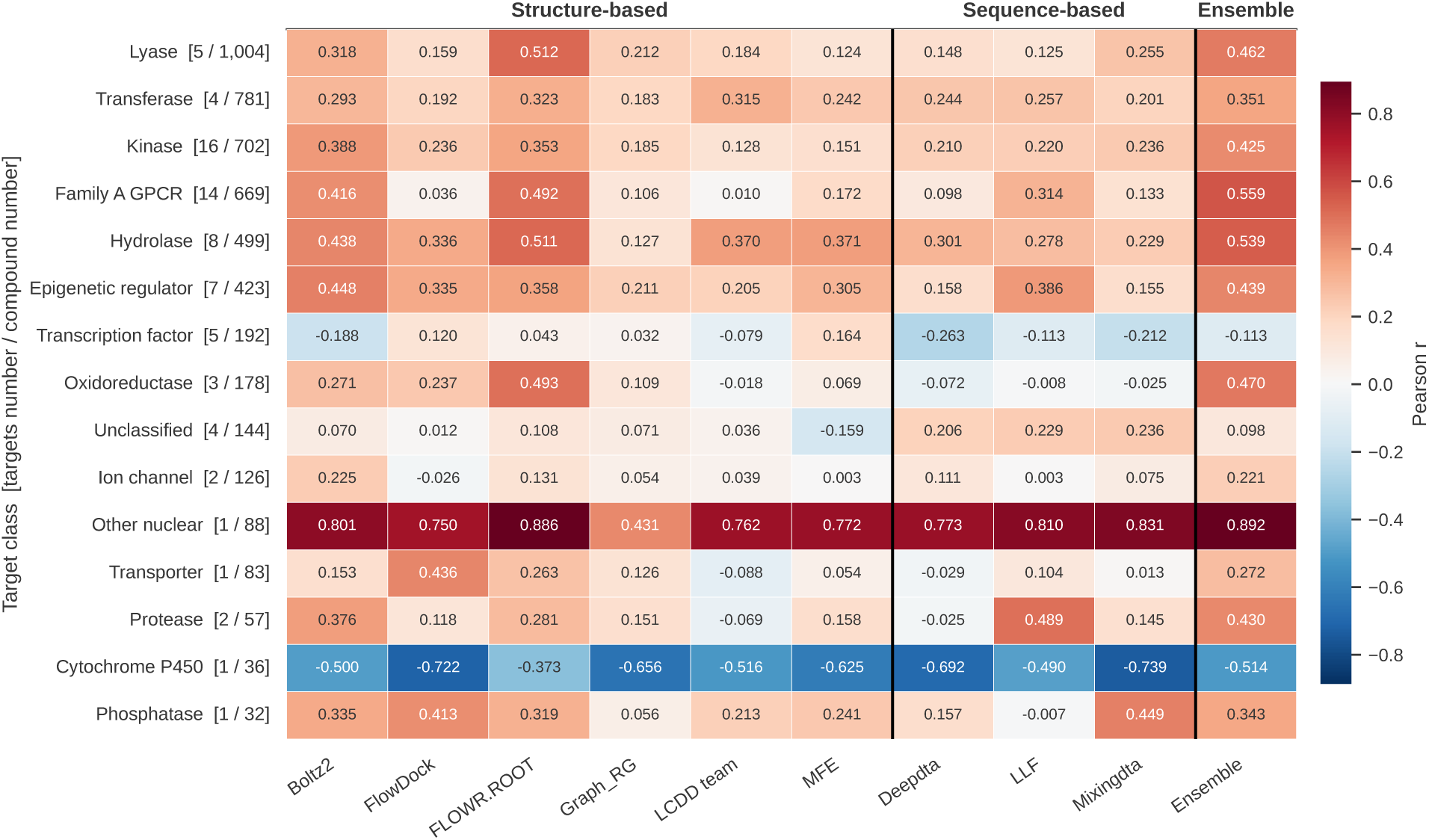
Class-conditional predictive performance across distinct protein families. Heatmap displays target-aggregated Pearson correlation coefficients (*r*) across 15 protein functional classes (*n* = 5,014 evaluation pairs across 73 unique targets from ChEMBL35 and CASP16; color scale ranges from *−*0.9 to 0.9, with red indicating positive linear correlation and blue indicating negative correlation). Target families along the *y*-axis are annotated as [number of targets, number of complexes]. Columns correspond to the nine individual deep learning models alongside a simple ensemble model. The simple ensemble is constructed by taking the unweighted arithmetic mean of binding affinity predictions from the two leading pretrained models, FLOWR.ROOT and Boltz2.

First, massively pre-trained models (e.g., FLOWR.ROOT, Boltz2) maintain a higher performance in large, well-characterized families such as Kinases, Hydrolases, and Lyases, indicating that scaling structural pre-training is a reliable method for learning standard pocket geometries.

Second, sequence-based methods consistently outperform structure-based approaches across all four targets in the Unclassified category. This superiority cannot be attributed to inadequate structural modeling: all four targets in the category possess experimental structures in the PDB with 100% sequence identity, and P17931 achieved an AF3 pLDDT of 97.8, the highest among all 71 targets. Instead, the performance gap is driven largely by NLRP3, which contributes nearly half of the complexes in this category (69 of 144 ligands) and exhibits a Kendall’s *τ* margin of 0.247 in favor of sequence-based models. Consequently, the availability of well-characterized protein templates does not ensure a performance advantage for structure-based scoring on these targets.

Third, nearly all evaluated models show markedly reduced performance on the transcription factor, Cytochrome P450, and ion-channel targets represented in our benchmark, suggesting potential shared limitations in modeling these complex biochemical contexts. For Cytochrome P450 on which all the models failed, enzymatic activity critically depends on the catalytic heme group. However, among the evaluated models, only LCDD-team and MFE explicitly support metal ions and organic cofactors (while FLOWR.ROOT supports only organic cofactors). Heme-containing complexes constitute merely *∼*0.1% of the PDBbind training corpus, rendering heme-mediated binding heavily out-of-distribution (OOD) even for the cofactor-aware architectures. Consequently, this failure reflects a limitation of existing models when applied to complex biochemical contexts.

Similarly, for ion channels, severe performance degradation is observed across almost all evaluated methods, raising the question of whether the failure stems from supplying only single-chain monomeric inputs to the models. To rigorously verify this, we reconstructed the homo-pentameric *α*7 nAChR (*n* = 19) and homo-tetrameric hERG (*n* = 20) multimer assemblies. Specifically, we used experimental crystal structures (PDB: 8V8A for *α*7 and 8ZYQ for hERG) as conditions for pose generation and selected ligand poses maximizing contact-residue fingerprint overlaps to approximate physiological binding conformations. However, presenting the complete assembly as input does not yield a systematic gain across absolute error, correlation, and ranking metrics. For *α*7, while multi-chain inputs reduce absolute RMSE for four of the six models (except FLOWR.ROOT from 0.59 to 0.84), correlation and ranking capabilities remain essentially random. In contrast, for hERG, multimeric inputs considerably inflate RMSE (e.g., LCDD-team increasing from 1.24 to 2.36 and MFE from 1.45 to 2.22; paired bootstrap 95% confidence intervals exclude zero) without statistically meaningful ranking improvements.

These findings indicate that providing multimeric inputs does not resolve the predictive bottleneck on these two ion channel targets. This suggests that structural fidelity alone may not be the sole limiting factor in multimeric affinity prediction, consistent with our observations in Fig. 3 that eliminating structural inaccuracies does not necessarily yield superior affinity scoring. Two target series cannot establish a general deficiency, but the absence of any gain indicates that oligomeric state alone is not the primary limiting factor.

#### Complementary model behavior suggests opportunities for class-aware prediction

Fig. 4 shows that no single method achieves universal supremacy across all protein classes. For example, while Boltz2 is second in general, it is among the worst performers for Transcription factors (*r* = *−*0.188). Similarly, FlowDock surpasses the leading FLOWR.ROOT in Transporters (0.436) and Phosphatases (0.413), yet it performs randomly in Family A GPCRs (0.036) and Ion channels (*−*0.026). This protein class-dependent performance suggests that the current PLA prediction methods are complementary, and therefore it may be useful to combine them to further improve the prediction accuracy.

Beyond improving the precision of these critical targets, the ensemble may also serve as a pragmatic risk-mitigation tool against domain-specific failures. Motivated by this observation, we evaluated a straightforward ensemble strategy by averaging the predictions of the two leading models, FLOWR.ROOT and Boltz2. Globally, the simple ensemble maintains state-of-the-art performance, achieving an overall Pearson correlation of 0.410, higher than that of FLOWR.ROOT’s single-model peak (0.392). However, the most compelling advantage of this approach is observed in several protein classes that are highly relevant to drug discovery. The ensemble achieves the best performance across five major families: Transferases, Kinases, Family A GPCRs, Hydrolases and Other nuclear proteins, which cover *∼*54.6% of the test set. According to comprehensive pharmacological analyses,^32^ these specific families span the predominant classes of approved human therapeutic targets, led by GPCRs (*∼*30%) and major enzyme superfamilies (*∼*47%; primarily transferases including kinases, and hydrolases including proteases). In contrast, excluding Cytochrome P450 (*∼*0.7% of the benchmark), where all evaluated models failed to achieve positive correlation (FLOWR.ROOT *r* = *−*0.373), the two enzyme families where FLOWR.ROOT maintains a lead (Lyase and Oxidoreductase) account for 23.6% of the filtered dataset, but represent only *∼*8% of approved human drug targets.^32^ Therefore, in addition to achieving a strong overall performance, the ensemble strategy implicitly improves the predictive precision across high-value pharmacological target classes.

While ensembling can moderately average down the peak performance of individual models on certain targets (e.g., Epigenetic regulators and Oxidoreductases), it provides a practical mitigation strategy against domain-specific failures of individual models. For example, in the Lyase family, the largest target class in the benchmark, the ensemble draws on FLOWR.ROOT’s superior accuracy (*r* = 0.512 versus 0.318 for Boltz2) to maintain a robust correlation of *r* = 0.462. However, when both constituent models share the same blind spots, as in Transcription factors and Cytochrome P450, the ensemble cannot overcome these inherent limitations, yielding correlations of *−*0.113 and *−*0.514, respectively.

## Discussion

In this study, we introduce PLABench, a comprehensive and standardized evaluation framework, to address the critical limitations of data leakage and potential bias in evaluating the prediction of protein–ligand binding affinity (PLA). By bridging the gap between sequence-based and structure-based paradigms and incorporating high-accuracy AlphaFold3-predicted structures, PLABench provides a rigorous, target-centric assessment of the generalizability of current deep learning models for PLA prediction.

Our evaluations reveal a dual narrative in the current state of deep learning-driven drug discovery. On one hand, the performance advantage of FLOWR.ROOT and Boltz2 over other evaluated models is plausibly attributable to their substantially larger training datasets. However, because architectures, training objectives, and data compositions also differ, scale cannot be isolated as the sole cause. A common, large-scale, and high-quality training corpus would enable future work to separate algorithmic gains from data-scaling effects.

On the other hand, our evaluation reveals some fundamental vulnerabilities of the current models. As demonstrated on the CASP16 dataset, high regression accuracy frequently masks a severe inability to accurately rank congeneric ligands in low-variance, high-affinity regimes. This limitation is further exacerbated when models encounter unfamiliar chemical spaces or complex biological mechanisms. In particular, our target-centric analysis on ChEMBL35 reveals markedly reduced performance across the Cytochrome P450 and transcription-factor targets with all evaluated architectures. However, as our analysis indicates, the performance degradation on these targets may be due to multiple factors such as methodological shortcomings, limited training data, and current input-side constraints. Specifically, expanding architectural support and training coverage for essential cofactors (such as catalytic heme groups) remains an unfulfilled need.

To systematically examine how structural quality governs the behavior of structure-based methods, we evaluated the CASP16 L3000 series across inputs spanning a spectrum of conformational qualities. In particular, increasing structural accuracy does not translate into monotonic improvements in affinity prediction for advanced models. Even when supplied with ground-truth experimental crystal structures, several structure-based architectures fail to outperform a sequence-only baseline, MixingDTA. A consistent pattern emerges on ChEMBL35, where sequence-based models demonstrate superior robustness across multiple targets. These results suggest that structure-based methods need to better leverage binding-mode features to genuinely link 3D interactions with affinity.

Finally, PLABench reveals the protein-class-dependent performance of different PLA prediction methods on different protein families and their complementarity. Class-aware routing or ensembling of complementary models offers a pragmatic strategy for immediate performance gains. Closing the gap on cofactor-dependent and underrepresented targets will also require architectures that incorporate explicit binding contexts and expanded training corpora to cover these classes.

## Methods

### PLABench framework overview

PLABench (Fig. 1) evaluates protein–ligand affinity predictors under three conditions rarely unified in existing benchmarks: strict temporal isolation from training data, standardized structural inputs bridging sequence- and structure-based paradigms, and target-centric scoring. The framework reports both absolute prediction error and within-target ranking. Its automated pipeline handles data preprocessing, target grouping, and metric computation, enabling new models to be evaluated directly from standardized prediction outputs.

### Datasets

PLABench integrates diverse datasets to comprehensively evaluate models in varying data scenarios and bridge the modality gap between sequence- and structure-based paradigms. We include two classic sequence-based benchmarks, namely Davis and KIBA, to assess generalization capabilities. More importantly, to minimize data leakage and enable rigorous evaluation of generalization, we incorporate the newly published ChEMBL35 and CASP16 blind-test datasets, both of which provide high-quality, target-centric congeneric ligand series. To ensure a fair head-to-head comparison of sequence-based PLA prediction models, we utilize a unified training corpus based on the PDBbind v2020 Refined set, upon which all sequence-based methods in PLABench were retrained. Furthermore, we provide curated AlphaFold3^19^-generated structures for the targets in the CASP16 and ChEMBL35 datasets to fairly compare the structure-based PLA prediction methods. This inclusion also allows us to rigorously evaluate whether structure-dependent models can maintain robust predictive accuracy in realistic drug discovery settings where experimental crystal conformations are unavailable.

#### Davis dataset

The Davis dataset^26^ comprises binding affinity (p*K*_d_) data for kinase protein families and their corresponding inhibitors. To ensure a rigorous evaluation and prevent information leakage observed in previous studies^4, 33^ utilizing raw datasets, we systematically standardized the data. Duplicated protein–ligand pairs were removed, and conflicting affinity values for identical pairs were resolved by retaining the maximum recorded value, yielding a highly curated dataset of 25,772 unique protein–ligand interaction pairs across 379 unique protein sequences (originally 442 kinase entries, with isoforms sharing identical sequences merged) and 68 ligands.

Given the limited diversity of unique targets and ligands in this dataset, standard random data splits cannot rigorously test a model’s ability to generalize to new proteins or ligands not observed in training data. Therefore, in addition to the standard random partitioning (i.e., evaluated using consistent 5-fold cross-validation on an 80/20 train/test split to ensure an equitable baseline), we adopted the rigorous cold-target and cold-drug split strategies established by MixingDTA.^5^ By strictly including only unseen targets or unseen chemical structures into the test set, this approach aligns with recent state-of-the-art evaluation protocols, enabling a strict assessment of a model’s generalizability to unseen biological and chemical spaces.

#### KIBA dataset

The KIBA dataset^27^ was constructed to address data heterogeneity in public bioactivity databases by integrating complementary information from multiple assay metrics (IC_50_, *K*_i_, and *K*_d_) into a single unified KIBA score. This integration approach yields a highly consistent and robust benchmark for comparing kinase-inhibitor bioactivity across diverse experimental settings.

Consistent with our evaluation protocol and the standards established by MixingDTA,^5^ we utilized the Therapeutics Data Commons (TDC)^34^ version of the KIBA dataset in PLABench. This highly curated set comprises 117,657 drug–target interaction pairs, covering 229 proteins and 2,068 ligands. Following the identical standardization protocol applied to the Davis dataset, we evaluated models across three rigorous partitioning strategies: a random split, a cold-target split, and a cold-drug split.

To resolve the inherent discrepancy between standard cross-validation and strict cold-splitting, we adopted a two-stage evaluation methodology for the cold datasets. Taking the cold-drug split as an example, we first held out approximately 20% of the unique drugs to construct a fixed, completely unseen test set. A 5-fold cross-validation scheme was then strictly confined to the remaining data (maintaining a 4:1 training-to-validation ratio per fold). Ultimately, the five models derived from all five folds were evaluated on the same fixed cold-drug test set separately, and their final predictions were aggregated. The cold-target split followed an analogous protocol.

#### PDBbind v2020 dataset

The PDBbind database serves as the gold-standard corpus for structure-based drug design, systematically pairing experimental 3D biomolecular structures from the Protein Data Bank (PDB)^35^ with their corresponding high-quality binding affinity data. To enable a fair comparison between sequence-based and structure-based models on the subsequent independent test sets (i.e., CASP16 and ChEMBL35), we retrained the sequence-based models using the same PDBbind v2020 refined set,^25^ following the training and evaluation protocol established in MixingDTA.^5^ Notably, our setup introduces two distinct modifications: rather than directly adopting the published input data from MixingDTA, protein sequences and ligand representations were reconstructed from the raw structural data; furthermore, we replaced their five-fold warm-start training scheme with a randomized 90/10 split, holding out the 10% partition as a validation set for early stopping.

Regarding data curation, from the initial 5,316 entries in the PDBbind refined set, we excluded 342 complexes overlapping with the CASF-2013^9^/CASF-2016^10^ core sets or CSAR-HiQ 36/51^11^ benchmarks, as well as an additional 12 entries with censored (*≤*) affinity annotations, missing affinity records, or unparseable ligands, yielding a final set of 4,962 complexes.

Some other PDBbind preprocessing pipelines^5, 36^ construct target sequences by concatenating all resolved chains end-to-end and truncating the concatenated string. In contrast, to ensure consistency with our benchmarks that inherently provide only single-chain sequences, such as CASP16 and ChEMBL35, we standardized our pipeline on the longest resolved chain when training and testing sequence-based models. To directly quantify the impact of this choice, we conducted an ablation study retraining all three sequence-based models on concatenated multi-chain inputs versus longest single-chain representations. Across three random seeds evaluated on six test sets, switching between representations shifted the mean Pearson correlation by only 0.016. This difference is less than half the standard deviation of 0.033 induced purely by altering random initialization seeds, with metric variations showing no systematic bias toward either preprocessing strategy ( Supplementary Note 1, Supplementary Tables 1 and 2). Accordingly, we report and discuss sequence-based models trained on the longest single-chain representation throughout the paper.

#### CASP16 dataset

The 2024 CASP16 competition^28^ introduced a dedicated protein–ligand binding affinity prediction category, establishing a community-wide blind benchmark for evaluating computational scoring functions. Sourced directly from active pharmaceutical drug discovery research, the challenge comprises two clinically relevant targets with congeneric ligand series: Chymase (series L1000, *n* = 17 ligands) and Autotaxin (series L3000, *n* = 123 ligands). The bioactivity data reflect highly relevant clinical scenarios, with affinities ranging from 1 nM to 10 µM and experimental crystal resolutions between 1.1 and 2.7 Å.

Unlike standard benchmark datasets characterized by broad and diverse affinity distributions, the CASP16 series is distinguished by its concentrated, high-affinity regimes with small variance. Consequently, standard global error metrics are insufficient; rather, the official assessment strictly emphasizes the ability of the models to accurately capture the relative ranking of congeneric ligands within these narrow windows, utilizing Kendall’s *τ* as the primary metric. Crucially, to integrate this rigorous challenge into PLABench and support structure-based methods under realistic blind-test conditions, we provide standardized high-accuracy AlphaFold3-predicted structures for all the target complexes, enabling a fair evaluation of model performance when experimental conformations are unavailable. Specifically, we generated 10 poses for each protein–ligand complex and selected the optimal conformation based on the highest ipTM score. Similarly, Boltz2^6^ was configured to sample 10 structural conformations and evaluate binding affinity in the highest-ipTM pose with 500 sampling steps and 5 recycling steps to reproduce the performance reported on the CASP16 Dataset.

#### ChEMBL35 dataset

The ChEMBL35 dataset^31^ was curated to serve as a leakage-controlled external benchmark for assessing model generalizability. Based on the curation protocol of Shimizu et al.,^29^ we further discarded any test ligand with a maximum similarity of ECFP4 Tanimoto *≥* 0.85 to any training molecule, regardless of protein target, thus eliminating both exact matches and close structural analogs.

Furthermore, we applied rigorous biochemical filters to align the test set with the intrinsic predictive capabilities of current scoring functions. Specifically, we excluded data points annotated with EC_50_ values, because EC_50_ conflates binding affinity with downstream signal amplification and is therefore not directly comparable to *K*_d_, *K*_i_, or IC_50_ on the same affinity scale. Moreover, the standard PDBbind training set contains a negligible fraction (approximately 0.14%) of EC_50_ records, rendering models trained on it ill-equipped for such predictions.

Finally, to facilitate robust target-centric ranking analyses, we removed any protein targets associated with fewer than 10 ligands. The final curated dataset comprises 71 unique protein sequences paired with 4,862 ligands. Crucially, consistent with our CASP16 protocol, we generated and publicly released high-confidence AlphaFold3 structures for all protein–ligand complexes within this dataset, enabling standardized, head-to-head evaluations of structure-based methods without relying on experimental crystal conformations.

Following the benchmark curation of Shimizu et al.,^29^ all target structures were evaluated as single-chain monomers. Specifically, their curation pipeline selected targets based on ChEMBL’s “SINGLE PROTEIN” classification. In ChEMBL data conventions, this definition indicates only that a target is encoded by a single gene (a unique UniProt accession); while this effectively excludes heteromeric complexes, it inadvertently retains homomultimeric targets in single-chain formats. Consequently, for targets whose functional forms are homomultimeric (e.g., the homo-pentameric nAChR *α*7 and homo-tetrameric hERG), monomeric representations inevitably omit cross-subunit interfaces where ligands bridge adjacent chains. While resolving physiological multimer assemblies and validating their binding pockets across the entire benchmark remains an essential direction for future dataset iterations, we explore the impact of this factor through targeted multimer modeling (see Results, ‘Large-scale pretraining improves overall generalization but does not eliminate target-dependent failures’).

### Prediction tasks

To comprehensively evaluate the utility of deep learning models across different stages of drug screening, PLABench frames the protein–ligand binding affinity prediction challenge into two distinct but complementary computational tasks below.

#### Absolute affinity Prediction (global regression)

The objective of this task is to map a given protein–ligand pair to a continuous scalar value that represents its absolute thermodynamic binding strength (e.g., p*K*_d_, p*K*_i_ or a unified KIBA score). Accurately predicting global affinity helps select ligands with strong binding affinity in early stage drug discovery.

#### Target-centric relative ligand ranking

This task evaluates a model’s capacity to accurately order a congeneric series of ligands against a single, specific protein target based on their relative binding strengths. As demonstrated in our evaluations, models that achieve low global regression errors may fail at this task, especially within high-affinity, low-variance regimes. Success in this task requires models to capture subtle stereochemical variations and NCIs among ligands rather than predicting mean binding affinity. This task is important for ranking ligands during lead optimization.

### Evaluation metrics

To comprehensively evaluate the models across the two predefined computational tasks, global regression and target-centric relative ranking, we selected seven representative metrics. Throughout this section, let *N* represent the total number of test samples (target–ligand pairs), *y_i_* denote the experimental (true) binding affinity for the *i*-th sample, and *ŷ_i_* denote the predicted binding affinity.

#### Mean squared error

The mean squared error (MSE) is the average of the squared differences between predicted values and true values, quantifying the overall magnitude of prediction errors. MSE is defined in Equation (1).

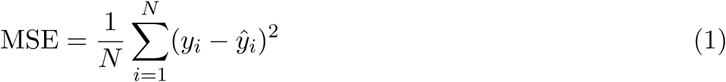

#### Root mean squared error

The root mean squared error (RMSE) is the square root of the MSE, as defined in Equation (2).

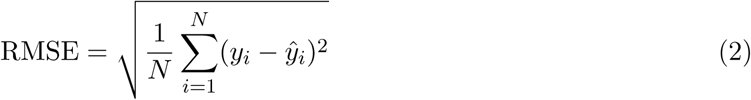

#### Pearson correlation coefficient

The Pearson correlation coefficient (PCC) is a classic metric for measuring the strength and direction of a linear relationship between two continuous variables. Ranging from -1 to 1, this coefficient is derived from the ratio of the covariance of the true and predicted values to the product of their respective standard deviations, mathematically defined in Equation (3). Here, *ȳ* and *ŷ̄* represent the mean of the true and predicted values, respectively. A value of 1 indicates a perfect positive correlation, -1 denotes a perfect negative correlation, and 0 suggests no linear association.

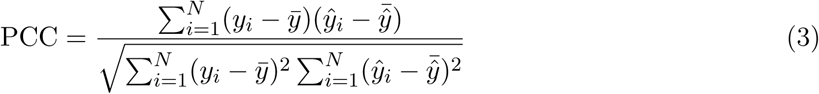

#### Spearman’s rank correlation coefficient

Spearman’s rank correlation coefficient (*ρ*) assesses the monotonic relationship between predicted and true values by converting them into ranks, rather than relying on the original continuous data. It is robust to non-normal data distributions and outliers, making it highly suitable for evaluating non-linear monotonic correlations in PLA prediction. The metric is defined in Equation (4), where *d_i_* = *R*(*y_i_*) *− R*(*ŷ_i_*) represents the difference in ranks between the true and predicted values for the *i*-th sample. Because tied values occur in both experimental and predicted affinities, *ρ* is calculated as the Pearson correlation between the average ranks of *y* and *ŷ*, which reduces to Equation (4) in the absence of ties.

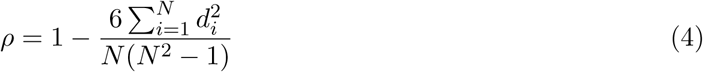

#### Modified squared correlation coefficient

The modified squared correlation coefficient (*r*^2^_m_), introduced by Roy et al.,^37^ penalizes predictions that correlate with the experimental values but are systematically offset from them:

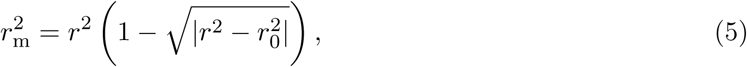

where *r*^2^ is the squared Pearson correlation between predicted and experimental affinities, and *r*^2^_0_ is the coefficient of determination of the regression of experimental on predicted values through the origin. Specifically, *r*^2^_m_ reduces to *r*^2^ only when the linear regression of experimental on predicted values passes through the origin; it penalizes non-zero intercepts (additive offsets) rather than pure scale errors.

#### Concordance index (CI)

The concordance index (CI) evaluates the global ranking performance of a model, defined as the probability that the predicted order matches the true order for a randomly selected pair of comparable samples:

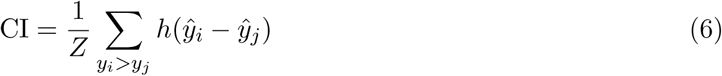

where *Z* = ∑_*y_i_ > y_j_*_ 1 is a normalization constant representing the total number of valid pairs with distinct experimental values (*y_i_ > y_j_*), and *h*(*u*) is a step function that rewards concordant predictions and assigns half credit to tied predictions:

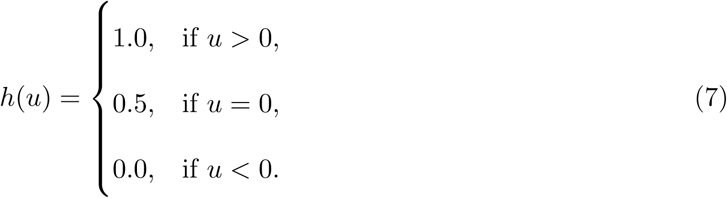

The CI value ranges from 0 to 1, where 0.5 corresponds to random guessing and 1.0 denotes perfect rank concordance.

#### Kendall’s *τ*

Kendall’s *τ* measures the strength of rank correlation between two variables based on concordant and discordant pairs. For all possible sample pairs, if the ordering direction of the true and predicted values is consistent, it is a concordant pair (*C*); otherwise, it is a discordant pair (*D*). The calculation of *τ* is expressed as the normalized ratio of the difference between concordant and discordant pairs to the total number of pairs, defined in Equation (8). Here, *T_y_* and *T_y_*_^_ denote the number of ties in the true and predicted values, respectively. Ranging from -1 to 1, a higher *τ* indicates stronger agreement. We utilized this metric extensively for the CASP16 and ChEMBL35 datasets, as it is the official evaluation standard designated by the CASP16 organizers and is exceptionally well-suited for rigorously assessing a model’s relative ranking capabilities across congeneric ligands targeting the same protein.

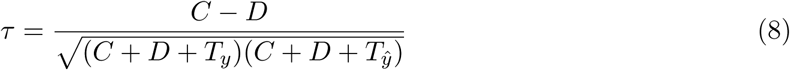

## Data availability

The data generated in this study have been deposited in Zenodo under accession code doi:10.5281/zenodo (ref.^38^). These comprise: (1) benchmark inputs and split partitions (PDBbind v2020 refined partition of 4,465 training and 497 validationcomplexes, given as PDB codes and split labels; the Davis and KIBA warm, cold-target and cold-drug fold assignments; the filtered ChEMBL35 test set of 4,862 ligand–target pairs across 71 targets with protein-family annotations, alongside the 7,650-pair unfiltered set and the per-target removal ledger; the CASP16 L1000 and L3000 targets converted to p*K*_d_; and the standardized CSAR-HiQ 36 and 51 tables); (2) AlphaFold3-predicted structures for the ChEMBL35 and CASP16 test sets; (3) the retrained sequence-model and MFE checkpoints; and (4) all per-model predictions, per-ligand leakage-audit tables and evaluation metrics. A provenance table in the deposit (SOURCES.tsv) gives the origin and license of every file. The AlphaFold3-predicted structures are distributed under the AlphaFold3 Output Terms of Use for non-commercial use, with the conversion from mmCIF to separated protein and ligand files; no AlphaFold3 model parameters are redistributed. The ChEMBL35 files derive from ChEMBL release 35 and are deposited under CC BY-SA.

The following third-party datasets were analyzed but are not redistributed here, and are available from their original sources: the PDBbind v2020 refined and general sets (http://www.pdbbind.org.cn/, mirrored at https://huggingface.co/datasets/photonmz/pdbbindpp-2020), whose licence does not permit redistribution; the CASF-2013 and CASF-2016 core sets (http://www.pdbbind.org.cn/casf.php); the Davis and KIBA datasets via Therapeutics Data Commons (https://tdcommons.ai/); the CASP16 ligand-affinity targets L1000 and L3000 (https://predictioncenter.org/casp16/); ChEMBL release 35 (https://ftp.ebi.ac.uk/pub/databases/chembl/ChEMBLdb/releases/chembl_35/); the curated ChEMBL35 benchmark from Shimizu et al.^29^ (https://zenodo.org/records/18669539); the standardized CSAR-HiQ 36 and 51 affinity tables,^11^ whose portal has been discontinued and which remain mirrored at https://bindingmoad.org/; and the Protein Data Bank entries 8V8A and 8ZYQ (https://www.rcsb.org/).

Source data for Figs. 2–4, Table 2-4 and Supplementary Tables 1–4 are provided with this paper.

## Code availability

The full implementation, evaluation pipelines and reproduction scripts are available at https://github.com/BioinfoMachineLearning/PLABench under an MIT license. The version used in this study is archived on Zenodo under https://doi.org/10.5281/zenodo.22817205.39 Versions and commit hashes of all third-party models and tools are listed in the README of the repository. Third-party model weights are not redistributed and are fetched from their original releases with checksum verification by scripts/download\_third\_party.sh.

## Acknowledgements

We thank all members of the Bioinformatics and Machine Learning Laboratory (BML) at the University of Missouri for constructive discussions and feedback, with special thanks to Dr. Alex Morehead and Pawan Neupane for their valuable insights. Computational resources were supported by the University of Missouri.

## Author contributions

J.C. conceived the study. L.W. and J.C. designed the experiments. L.W. curated the benchmark datasets, generated AlphaFold3 complex structures, developed the evaluation pipelines, performed computational experiments, collected the experimental data, and conducted the statistical analyses. J.C. supervised the research and guided the investigation. L.W. and J.C. analyzed the results and wrote the manuscript.

