## Supplementary Information for "Leakage-controlled benchmarking reveals generalization limits of deep learning for protein–ligand binding affinity prediction"

---

### Supplementary Notes

#### Supplementary Note 1 Target chain-selection ablation for sequence-based models

##### Experimental design

Because sequence-based binding affinity predictors accept a single contiguous amino acid sequence per target, benchmark pipelines derived from the Protein Data Bank (PDB) can define an explicit rule when handling multimeric complexes. We selected the longest resolved chain as the representative sequence, whereas several published studies concatenate all resolved chains end-to-end.[1,2] Among our benchmark suites, only the PDBbind-derived sets (CASF-2013, CASF-2016, CSAR-HiQ 36, and CSAR-HiQ 51) contain multichain entries requiring chain selection; CASP16 and ChEMBL35 provide a single canonical sequence per target. Consequently, identical input representations across CASP16 and ChEMBL35 ensure that any performance differences arise solely from the retrained model weights.

To evaluate potential artifacts introduced by this convention, each sequence-based model (LLF, DeepDTA, and MixingDTA) was retrained on the PDBbind v2020 refined set (4,465 training and 497 validation complexes) under both protocols; for a given model, the longest-chain and concatenated strategies used the same hyperparameters, optimizer, and early-stopping criterion, so that the two settings differed only in the target sequence representation. Both settings employed a uniform 4,700-token sequence window to avoid confounding chain selection with sequence truncation. Evaluations were conducted via a standardized, automated scoring harness across three random seeds for LLF and DeepDTA, and a single run for MixingDTA due to its much larger computation amount.

##### Effect of the chain-selection rule

As summarized in Supplementary Table 1, performance differences between concatenating all chains and retaining the longest chain are negligible across all six test sets. Resampling variability was quantified using paired bootstrapping matching the reporting unit of each dataset (complexes for PDBbind sets, series-level rows for CASP16, and targets for ChEMBL35).

Across models trained across multiple seeds, the background variance across random initialization seeds (SD = 0.033) exceeded the mean absolute shift attributable to the chain-selection rule

($|\Delta| = 0.016$) by approximately two-fold. Only five of fifty-four metric comparisons reached nominal significance ($p < 0.05$), and none exceeded the variance observed across random seeds on the corresponding test set (e.g., $|\Delta| = 0.081$ versus seed standard deviations of 0.082–0.111 for LLF on CSAR-HiQ 36). A parallel evaluation on RMSE yielded identical conclusions, confirming that the chain-selection convention does not introduce systematic performance bias.

#### Difference-in-differences analysis on multichain targets

To determine whether sequence concatenation imparts a genuine biophysical advantage on multimeric proteins, we conducted a difference-in-differences (DiD) analysis comparing multichain complexes against single-chain controls (Supplementary Table 2). Because single-chain complexes receive identical sequences under both conventions, a true representational benefit from concatenation must yield a positive interaction term ($\text{DiD} = \Delta_{\text{multi}} - \Delta_{\text{single}} > 0$).

Across all evaluated models, no positive DiD estimate reached statistical significance. The sole significant effect was observed for DeepDTA on CASF-2016 ($\text{DiD} = -0.068$, $p < 0.001$), but its negative sign reveals that concatenation yielded greater nominal improvements on single-chain complexes whose inputs were unchanged. This demonstrates that performance variations stem from retraining stochasticity rather than the inclusion of multichain sequence content.

#### Structural and mechanistic interpretation

These empirical observations align with the architectural constraints of 1D sequence encoders. Because standard 1D CNNs, GCNs, and sequence transformers omit boundary tokens distinguishing subunit transitions, concatenated inputs are processed as continuous artificial polypeptides. The resulting residue representations undergo global adaptive pooling, destroying spatial and quaternary interface information. Retaining the longest resolved chain yields statistically equivalent performance while mitigating sequence truncation and false juxtaposition artifacts; we therefore adopted the longest-chain representation for all sequence-based models in PLABench.

**Supplementary Table 1 | Comparative performance of sequence-based models under concatenated versus longest-chain conventions across six benchmark datasets.**

| Model | Test set ($n$) | Metric | Longest | Concat | Δ | (95% CI) |
|---|---|---|---|---|---|---|
| LLF | CASF-2016 (285) | Pearson | 0.755 | 0.760 | +0.005 | (−0.011, 0.021) |
| | | Spearman | 0.735 | 0.744 | +0.009 | (−0.011, 0.028) |
| | | CI | 0.774 | 0.778 | +0.004 | (−0.005, 0.013) |
| | CASF-2013 (195) | Pearson | 0.647 | 0.660 | +0.013 | (−0.008, 0.034) |
| | | Spearman | 0.617 | 0.636 | +0.020 | (−0.008, 0.048) |
| | | CI | 0.722 | 0.731 | +0.008 | (−0.004, 0.021) |
| | CSAR-HiQ 36 (36) | Pearson | 0.467 | 0.548 | +0.081* | (0.015, 0.160) |
| | | Spearman | 0.461 | 0.521 | +0.061 | (−0.014, 0.155) |
| | | CI | 0.652 | 0.679 | +0.027 | (−0.013, 0.071) |
| | CSAR-HiQ 51 (51) | Pearson | 0.785 | 0.783 | −0.002 | (−0.045, 0.040) |
| | | Spearman | 0.701 | 0.683 | −0.018 | (−0.094, 0.047) |
| | | CI | 0.764 | 0.747 | −0.016 | (−0.053, 0.015) |
| | CASP16 (140) | Pearson | 0.444 | 0.412 | −0.031* | (−0.055, −0.007) |
| | | Spearman | 0.465 | 0.434 | −0.031 | (−0.068, 0.002) |
| | | CI | 0.665 | 0.653 | −0.012 | (−0.027, 0.002) |
| | ChEMBL35 (7650) | Pearson | 0.176 | 0.182 | +0.006 | (−0.004, 0.016) |
| | | Spearman | 0.165 | 0.170 | +0.005 | (−0.005, 0.015) |
| | | CI | 0.556 | 0.558 | +0.001 | (−0.002, 0.005) |
| DeepDTA | CASF-2016 (285) | Pearson | 0.746 | 0.745 | −0.001 | (−0.021, 0.020) |
| | | Spearman | 0.730 | 0.738 | +0.008 | (−0.018, 0.035) |
| | | CI | 0.770 | 0.771 | +0.001 | (−0.011, 0.012) |
| | CASF-2013 (195) | Pearson | 0.663 | 0.676 | +0.013 | (−0.013, 0.040) |
| | | Spearman | 0.649 | 0.663 | +0.013 | (−0.021, 0.048) |
| | | CI | 0.734 | 0.737 | +0.003 | (−0.012, 0.017) |
| | CSAR-HiQ 36 (36) | Pearson | 0.703 | 0.702 | −0.001 | (−0.076, 0.070) |
| | | Spearman | 0.739 | 0.729 | −0.010 | (−0.104, 0.087) |
| | | CI | 0.762 | 0.760 | −0.002 | (−0.052, 0.051) |
| | CSAR-HiQ 51 (51) | Pearson | 0.753 | 0.763 | +0.010 | (−0.038, 0.065) |
| | | Spearman | 0.654 | 0.706 | +0.052 | (−0.038, 0.148) |
| | | CI | 0.739 | 0.759 | +0.020 | (−0.024, 0.066) |
| | CASP16 (140) | Pearson | 0.469 | 0.497 | +0.028 | (−0.034, 0.080) |
| | | Spearman | 0.484 | 0.484 | +0.000 | (−0.068, 0.065) |
| | | CI | 0.664 | 0.667 | +0.003 | (−0.024, 0.029) |
| | ChEMBL35 (7650) | Pearson | 0.148 | 0.149 | +0.001 | (−0.026, 0.026) |
| | | Spearman | 0.143 | 0.142 | −0.001 | (−0.030, 0.023) |
| | | CI | 0.549 | 0.549 | −0.000 | (−0.010, 0.008) |
| MixingDTA | CASF-2016 (285) | Pearson | 0.781 | 0.788 | +0.007 | (−0.006, 0.021) |
| | | Spearman | 0.766 | 0.779 | +0.013 | (−0.005, 0.033) |
| | | CI | 0.790 | 0.795 | +0.005 | (−0.004, 0.014) |
| | CASF-2013 (195) | Pearson | 0.748 | 0.771 | +0.022* | (0.003, 0.044) |
| | | Spearman | 0.733 | 0.766 | +0.033* | (0.007, 0.064) |
| | | CI | 0.771 | 0.787 | +0.016* | (0.004, 0.031) |
| | CSAR-HiQ 36 (36) | Pearson | 0.700 | 0.712 | +0.012 | (−0.057, 0.079) |
| | | Spearman | 0.700 | 0.739 | +0.039 | (−0.047, 0.137) |
| | | CI | 0.762 | 0.779 | +0.017 | (−0.031, 0.065) |
| | CSAR-HiQ 51 (51) | Pearson | 0.732 | 0.731 | −0.000 | (−0.046, 0.045) |
| | | Spearman | 0.704 | 0.721 | +0.017 | (−0.070, 0.100) |
| | | CI | 0.758 | 0.774 | +0.016 | (−0.017, 0.052) |
| | CASP16 (140) | Pearson | 0.514 | 0.539 | +0.025 | (−0.002, 0.052) |
| | | Spearman | 0.529 | 0.552 | +0.023 | (−0.020, 0.068) |
| | | CI | 0.687 | 0.697 | +0.010 | (−0.007, 0.029) |
| | ChEMBL35 (7650) | Pearson | 0.145 | 0.144 | −0.002 | (−0.015, 0.013) |
| | | Spearman | 0.135 | 0.131 | −0.004 | (−0.017, 0.009) |
| | | CI | 0.547 | 0.546 | −0.001 | (−0.006, 0.003) |

Differences (Δ) denote concatenated minus longest-chain values; positive values indicate superior performance for concatenation. Asterisks (*) indicate where the 95% paired bootstrap confidence interval excludes zero. CASP16 metrics represent size-weighted aggregates over L1000 ($n = 17$) and L3000 ($n = 123$). ChEMBL35 is calculated per target across the 84 targets bearing $\geq 3$ compounds and weighted by target size. LLF and DeepDTA reflect 3-seed ensembles; MixingDTA is evaluated from a single model per setting.

**Supplementary Table 2 | Difference-in-differences (DiD) in Pearson correlation between multichain and single-chain strata.**

| Model | Test set | $n_{\text{multi}}$ | $n_{\text{single}}$ | $\Delta_{\text{multi}}$ | $\Delta_{\text{single}}$ | DiD | (95% CI) | $p$ |
|---|---|---|---|---|---|---|---|---|
| LLF | CASF-2016 | 98 | 187 | +0.011 | +0.002 | +0.010 | (−0.026, 0.046) | 0.572 |
| LLF | CASF-2013 | 76 | 119 | +0.023 | +0.004 | +0.019 | (−0.023, 0.065) | 0.369 |
| DeepDTA | CASF-2016 | 98 | 187 | −0.043 | +0.025 | −0.068* | (−0.114, −0.028) | <0.001 |
| DeepDTA | CASF-2013 | 76 | 119 | −0.011 | +0.035 | −0.047 | (−0.103, 0.005) | 0.082 |
| MixingDTA | CASF-2016 | 98 | 187 | +0.013 | +0.002 | +0.011 | (−0.020, 0.044) | 0.480 |
| MixingDTA | CASF-2013 | 76 | 119 | +0.032 | +0.013 | +0.019 | (−0.020, 0.063) | 0.360 |

$\Delta_{\text{multi}}$ and $\Delta_{\text{single}}$ indicate the Pearson correlation difference (concatenation minus longest chain) within multichain and single-chain subsets, respectively. Asterisk (*) denotes statistical significance at $p < 0.05$.

### Supplementary Note 2 Validation against official CASP16 competition submissions

Three structure-based architectures evaluated in PLABench were originally entered in the CASP16 blind challenge by their respective developers: Graph_RG (team Haiping, Group 16), LCDD-team (entered under the official competition name **BA-Pred**, Group 55; referenced as LCDD-team in the main text), and FlowDock (team MULTICOM_ligand, Group 207). In the official challenge, these teams generated bespoke 3D protein–ligand conformations and applied custom ranking protocols.

**Supplementary Table 3 | Concordance between official CASP16 submissions and standardized PLABench implementations.**

| Method | CASP16 group ID | Official CASP16 $\kappa_N$ | PLABench AF3 $\kappa_N$ | Difference [95% CI] |
|---|---|---|---|---|
| Graph_RG | Haiping (16) | 0.423 | 0.420 | −0.003 [−0.036, +0.029] |
| LCDD-team | BA-Pred (55) | 0.363 | 0.394 | +0.032 [−0.078, +0.141] |
| FlowDock | MULTICOM_ligand (207) | 0.319 | 0.319 | −0.001 [−0.104, +0.103] |

Evaluations are based on all 140 prospective blind targets (17 Chymase, 123 Autotaxin). $\kappa_N$ denotes the sample-size-weighted Kendall's $\tau$ adopted by official CASP16 assessors. Differences represent PLABench minus official submissions; 95% confidence intervals were derived from 5,000 paired bootstrap resamples.

To verify whether deploying standardized AlphaFold3 poses introduces negative distribution shifts, we retrieved the original prediction submissions from the CASP Prediction Center and scored them using our unified pipeline on the complete 140-compound dataset against blind experimental affinities. As shown in Supplementary Table 3, performance differences between official bespoke submissions and PLABench's standardized AlphaFold3 input pipeline are statistically indistinguishable ($|\Delta\kappa_N| \leq 0.032$; all 95% bootstrap confidence intervals encompass zero). This holds true even though MULTICOM_ligand utilized consensus scoring across multiple generative engines.

When subsetting to the 122 ligands evaluated in the official assessment report (excluding 18 ligands with prior disclosures), our re-scoring reproduced the official figures ($\kappa_N = 0.393$ versus 0.39 for Haiping, and 0.343 versus 0.33 for BA-Pred/LCDD-team). These results validate that off-the-shelf AlphaFold3 structures reliably represent targets in blind scoring evaluations without handicapping structure-based scoring heads.

### Supplementary Note 3 Sensitivity analysis of CASP16 rankings to training-set overlap

The CASP16 blind competition was conducted without filtering target–ligand pairs against the training corpora of deep learning predictors. To determine whether training memorization skewed model benchmarks, we performed an cross-corpus screen matching ligands by full canonical InChIKey against 14 bioactivity and structural databases.

Among the 123 Autotaxin (L3000) ligands, 26 were identified in historical databases, 21 of which possessed experimental binding affinities deposited in BindingDB between January 2021 and February 2022 (preceding the June 2023 pretraining cutoff). Twenty of the 21 appear, together with their labels, inside the training corpora of an evaluated model (19 in Boltz2's BindingDB and ChEMBL34 corpora and 14 in FLOWR.ROOT's SAIR and BindingNet corpora, with overlap between the two); the remaining ligand (L3021) is in neither corpus and was retained. For Chymase (L1000), only one ligand had an exact match. When enforcing strict triple-level overlap (identical target sequence, identical ligand InChIKey, and an attached affinity annotation), Boltz2 matched 19 Autotaxin complexes, and FLOWR.ROOT matched 14; the remaining seven methods exhibited zero training overlap.

**Supplementary Table 4 | Model performance on the de-duplicated CASP16 dataset excluding historical bioactivity overlaps.**

| | L1000 ($n = 16$) | | | | | | | L3000 ($n = 103$) | | | | | | |
|---|---|---|---|---|---|---|---|---|---|---|---|---|---|---|
| Model | MSE ↓ | RMSE ↓ | Pearson ↑ | Spearman ↑ | Kendall ↑ | $r^2_m$ ↑ | CI ↑ | MSE ↓ | RMSE ↓ | Pearson ↑ | Spearman ↑ | Kendall ↑ | $r^2_m$ ↑ | CI ↑ |
| Boltz2 | **0.447** | **0.669** | **0.546** | 0.391 | 0.233 | **0.077** | 0.617 | 1.139 | 1.067 | **0.596** | **0.595** | **0.427** | 0.247 | **0.714** |
| FlowDock | 2.436 | 1.561 | 0.138 | 0.112 | 0.100 | -0.005 | 0.550 | 1.446 | 1.202 | 0.423 | 0.422 | 0.288 | 0.134 | 0.644 |
| FLOWR.ROOT | 1.372 | 1.172 | 0.221 | 0.159 | 0.100 | 0.030 | 0.550 | **0.783** | **0.885** | 0.565 | 0.558 | 0.398 | **0.300** | 0.700 |
| Graph_RG | 1.387 | 1.178 | 0.431 | 0.444 | **0.283** | -0.010 | **0.642** | 1.202 | 1.096 | 0.515 | 0.556 | 0.382 | 0.222 | 0.692 |
| LCDD-team | 1.208 | 1.099 | 0.073 | 0.062 | 0.017 | 0.001 | 0.508 | 0.914 | 0.956 | 0.526 | 0.522 | 0.365 | 0.223 | 0.683 |
| MFE | 1.512 | 1.230 | 0.331 | **0.479** | 0.267 | 0.039 | 0.633 | 1.454 | 1.206 | 0.538 | 0.470 | 0.330 | 0.284 | 0.665 |
| DeepDTA | 1.071 | 1.035 | -0.196 | 0.244 | 0.183 | -0.006 | 0.592 | 1.710 | 1.308 | 0.387 | 0.352 | 0.253 | 0.115 | 0.627 |
| LLF | 0.568 | 0.753 | 0.188 | 0.309 | 0.200 | 0.022 | 0.600 | 3.669 | 1.916 | 0.354 | 0.331 | 0.235 | 0.111 | 0.618 |
| MixingDTA | 3.053 | 1.747 | -0.080 | 0.144 | 0.100 | -0.002 | 0.550 | 0.919 | 0.959 | 0.549 | 0.530 | 0.368 | 0.252 | 0.685 |

Metrics reflect the subset obtained after removing all target–ligand complexes whose exact structures and bioactivity annotations existed in historical databases prior to the competition (L1000: 16 of 17 retained; L3000: 103 of 123 retained).

Filtering out overlapping ligands ($n = 16$ for L1000; $n = 103$ for L3000) and recomputing rankings moderately reduced Kendall's $\tau$ across all methods, but preserved the relative model hierarchy (Spearman rank correlation between original and de-duplicated ranks: $\rho = 0.95$; Supplementary Table 4). The mean performance decrement in Kendall's $\tau$ was virtually identical between models with training overlap ($\Delta\tau = -0.082$) and the seven models with strictly zero overlap ($\Delta\tau = -0.086$).

This across-the-board drop reflects statistical dynamic-range compression rather than memory recall: the excluded historical ligands occupied the high-potency tail of the binding spectrum ($\mathrm{p}K_\mathrm{d} = 8.14 \pm 0.68$, compared to $6.70 \pm 1.05$ for retained compounds). Truncating the most potent binders narrows the evaluation variance, imposing a uniform mathematical penalty on rank correlation metrics across all predictors. These findings confirm that while absolute CASP16 metrics are elevated by broad assay dynamic ranges, the comparative conclusions and model leaderboards in PLABench remain uncompromised.
